# Thinking like a structural biologist: A pocket-based 3D molecule generative model fueled by electron density

**DOI:** 10.1101/2022.06.11.495756

**Authors:** Lvwei Wang, Rong Bai, Xiaoxuan Shi, Wei Zhang, Yinuo Cui, Xiaoman Wang, Cheng Wang, Haoyu Chang, Yingsheng Zhang, Jielong Zhou, Wei Peng, Wenbiao Zhou, Bo Huang

**Affiliations:** Beijing StoneWise Technology Co Ltd., Haidian Street #15, Haidian District, Beijing 100080, China; Innovation Center for Pathogen Research, Guangzhou Laboratory, Guangzhou 510320, China

**Keywords:** experimental electron density, deep learning, generative model, de novo drug design, structure-based drug design, three-dimensional generation.

## Abstract

We report for the first time the use of experimental electron density (ED) as training data for the generation of drug-like three-dimensional molecules based on the structure of a target protein pocket. Similar to a structural biologist building molecules based on their ED, our model functions with two main components: a generative adversarial network (GAN) to generate the ligand ED in the input pocket and an ED interpretation module for molecule generation. The model was tested on three targets including kinase (HPK1), protease (Covid19-3CL), and nuclear receptor (VDR), and evaluated with a reference dataset composed of over 8,000 compounds that have their activities reported in the literature. The evaluation examined the chemical validity, chemical space distribution-based diversity, and similarity with reference active compounds concerning the molecular structure and pocket-binding mode. Our model can reproduce classical active compounds and can also generate novel molecules with similar binding modes as active compounds, making it a promising tool for library generation supporting high-throughput virtual screening. Our model is available as an online service to academic users via https://edmg.stonewise.cn/#/create.

## Introduction

Molecular generative models using the three-dimensional (3D) information of target pockets have garnered increasing attention in the field of de novo drug design^1–3^. The process of drug design is generally perceived as an inherently multi-constrained optimization process. The major constraints include complementarity between the ligand and protein, regarding multiple aspects, such as shape and non-covalent interactions (NCI), and requirements of the ligand itself, including synthesizability and low strain energy binding conformation. Therefore, efficiently maintaining these constraints during training of the molecular generative model is subject to intensive discussions. Some representative attempts include the following: 1) using autoregressive algorithms or introducing conditional tokens in the training process for models generating molecules in a sequential manner^4^; 2) leveraging generative adversarial networks (GAN) and reinforcement learning to reflect the desired bias in the output distribution^5, 6^; 3) employing Bayesian optimization for the search of appropriate regions in the latent space from trained models such as variational autoencoder (VAE)^7^. These approaches all possess delicately designed architecture and have accomplished great achievements by working in an end-to-end manner in the fields of machine translation, games, and image processing. However, when applied to molecule generation for drug design, additional challenges arise, including an increasing number of constraints and the lack of data. As a result, the application of end-to-end design that intends to satisfy all the constraints at one stroke by using massive amounts of training data is not as effective as in other fields. Therefore, researchers pursue solutions in two directions: 1) reducing the constraints by generating non-3D molecules, in which the molecules are represented as strings (i.e. SMILES) or graphs^8–10^; 2) expanding the dataset, such as by generating ligand-protein complex data using docking approaches^11^. Although these attempts are inspiring, a groundbreaking strategy that can pour additional experimental data in the models is still absent. The current data expansion approaches are based on calculations heavily relying on computer-aided drug design (CADD) theory. This promotes artificial intelligence (AI) models learning from well-established rules instead of “real world” information.

To find a solution to the challenges posed by complicated constraints and insufficient data, an appropriate molecule representation, which can simplify the introduction of massive amounts of experiment data to the training process and more comprehensively reflect the nature of constraints than the traditionally used molecular representations, is crucial. There are primarily two traditional 3D molecule representations used for the AI model: the node-edge based representation^12, 13^ and dot-cloud-based representation such as Gaussian filtering^14^ and van der Waals radius^15^. Compared to the node-edge based representation in which one atom is located at one voxel, the dot clouds are continuously smoothing and more compatible with convolutional neural networks (CNN). However, regarding reflection of the physical and chemical properties of a molecule, such as intermolecular noncovalent interactions, bond order, and electrostatics, none of these traditional representations are ideal. Therefore, multiple channels are required to complementarily describe bond orders, elements, and NCI-related properties, such as hydrogen bond donor or hydrogen bond acceptor. Unfortunately, using these molecular representations does not alleviate the lack of data but rather aggravates the problem by creating a data sparsity problem for some channels (e.g., Br, I for elements, and sigma hole for NCI). In contrast, electron density (ED) is ideal for representing both the physical and chemical properties of a molecule: the ED intensity isosurface describes the shape of a molecule; electron localization function (ELF)^16^ describes the bond properties; and ED topology, such as saddle points, indicates the NCI between molecules^17^. As a result, there is no need to involve multiple channels, thereby avoiding the data sparsity problem. Moreover, ED is also the form in which massive amounts of experimental data are stored: there are over 120,000 high quality ED maps accumulated globally in the Protein Data Bank (PDB) over the past 60 years. Only part of the information in these experimental EDs is used for the determination of atom coordinates, whereas other information reflecting NCI and solvent distribution remains untapped^17^.

Based on the forementioned reasons, we propose the use of ED as the molecule representation in this work. In so doing, we are able to train our model with experimental ED. In addition, the design of our model is inspired by the procedures performed by structural biologists. This includes building molecules in the pocket, where the shape and key atoms are first defined based on their EDs, and stereochemistry constraints are then applied to improve conformation. We trained our model to learn constraints and generate molecules in two major steps. Initially, a GAN was used to take the ED of a pocket as input to learn pocket-ligand complementarity and generate the ligand ED. Subsequently, an ED interpretation module powered by vector quantized variational autoencoder (VQ-VAE2) ^18^, PixelCNN^19^, and V-Net^20^ was used to learn constraints on ligand validity and interpret the generated ED into molecules. Experimental EDs were used to train the GAN, and quantum mechanics (QM) based computational ED as well as force field based molecular conformations were used to train the ED interpretation module. A rule-based fragment assembler was also designed as part of such a module to ensure the validity of the generated molecules. We evaluated our model using classical CADD indicators, such as quantitative estimate of drug-likeness (QED)^21^ and synthetic accessibility score (SAS)^22^, to test the molecular validity. Additionally, we employed a set of more intuitive indicators to test whether our model could generate novel active compounds: 1) the reproduction of classical active compounds; and 2) the generation of novel molecules sharing a similar binding mode with the classical active compounds. Regarding diversity, chemical space distribution was used for measurement.

## Results

### Model Design

The model architecture (Fig. 1) was designed under the principle of learning constraints step-by-step: less abundant experimental ED data were used to learn the complementarity between ligands and pockets, and relatively abundant computational data were used to learn the constraints on molecule validity. Specifically, ED was used as a molecular representation to convert the molecules from discrete symbols to a continuous numerical representation. Subsequently, a GAN (Fig. 1a) was designed for the generation of ligand ED (filler ED) and trained with 27,006 ligand-protein complex experimental ED data to learn how the ligand ED could best complement the pocket ED. However, such a generated filler ED may not exist as a valid molecule, necessitating the VQVAE2-PixelCNN-VNet component for interpretation of the filler ED to valid and diversified molecules. By reconstructing 40 million semiempirical quantum mechanical (QM) level EDs (details in Methods section), the VQ-VAE2 (Fig. 1b) was trained to learn two discrete latent spaces (top-level and bottom-level). Next, an autoregressive sampling process powered by PixelCNN was conducted in the latent spaces using generated ligand ED as a condition so that the complementarity learned by the GAN was retained. After conditional sampling, the filler ED was converted into a group of reconstructed EDs. Then, the V-Net component takes the reconstructed ED as input to determine how to position the atoms to best maintain the shape of such ED and the type of atom required to best reflect the strength of the ED. The output of the V-Net component is called the map skeleton. A fragment library and a rule-based fragment substitution method were applied on map skeletons to create the final molecules.

**Figure 1.**
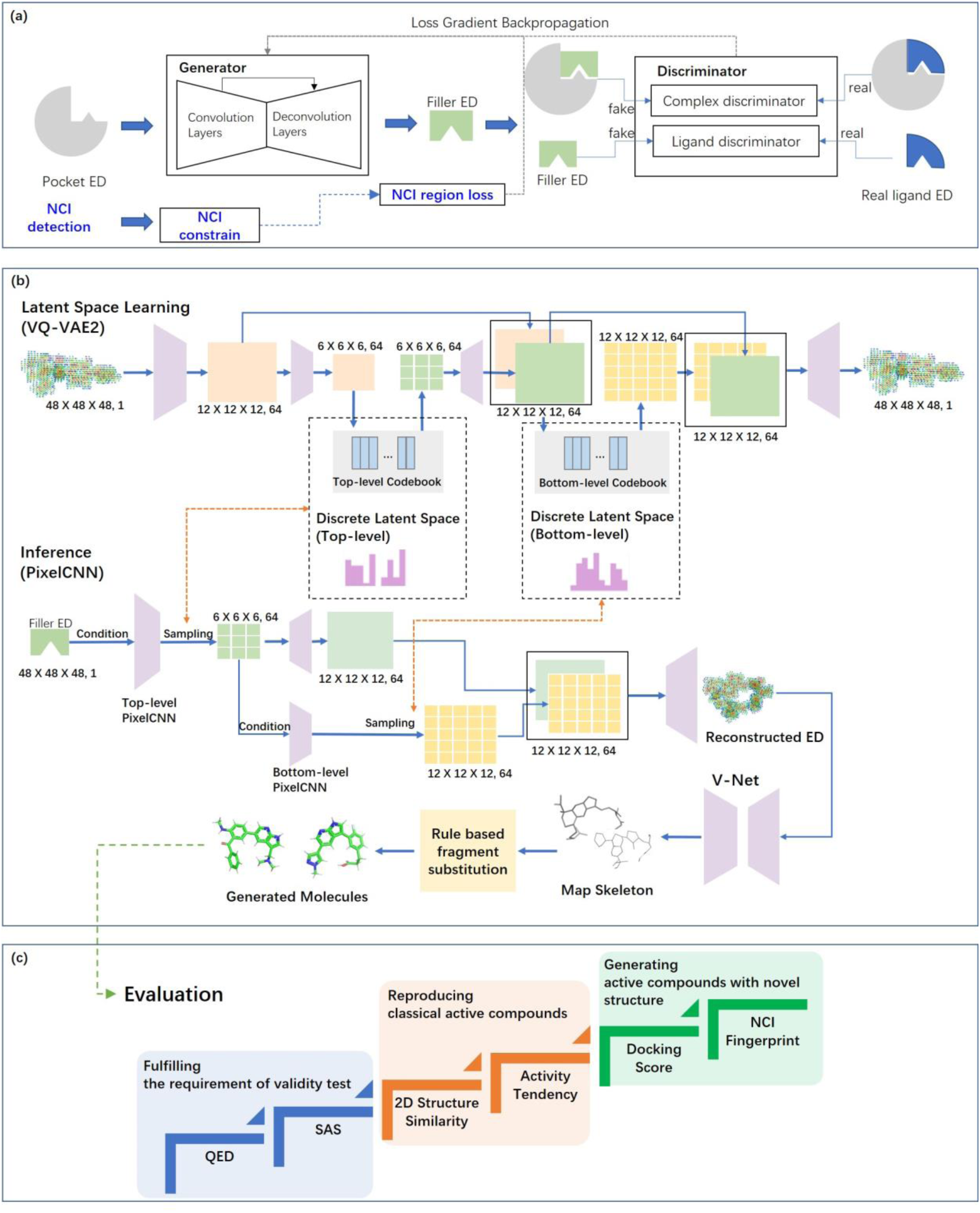
Model Architecture. a) The GAN for generating filler ED based on pocket ED. b) The training and inference process of VQVAE2-PixelCNN-VNet for generating molecules using filler ED as input. c) Illustration of evaluation framework. The blue part indicates the requirements for the validity test. The first level of the orange part indicates that the model can generate molecules that are previously reported active compounds; the second level indicates that the generated molecules should be more similar to high-activity compounds than low-activity compounds (i.e. a tendency to reproduce molecules with high activity). The green part represents the design meant to test whether there are active molecules with novel structures. All molecules with a low similarity (less than 0.5 Tanimoto similarity by using ECFP4) to the literature-reported active compounds were selected and first tested using the docking score and then tested using the binding mode by analyzing the NCI fingerprint. ED, electron density; GAN, generative adversarial network; NCI, non-covalent interactions; QED, quantitative estimate of drug-likeness; SAS, synthetic accessibility score; VQ-VAE, vector quantized variational autoencoder.

Benefitting from the fact that the ED intrinsically contains the information, such as shape, element, bond properties, and interactions, it is unnecessary to include extra channels in our model. To assure the efficiency of retaining constraints during training of the whole model, we increased the weight of the region where the ligand and pocket show non-covalent interactions (NCI) during the GAN training.

To demonstrate the functionality of our model, we applied the model to a kinase target, HPK1. The complex structure of HPK1 binding with a reference ligand (PDB:7KAC; Fig. 2a) was used as a starting point. The pocket was defined as all the residues within 5 Å from the reference ligand. Using experimental ED (2Fo-Fc map contoured at 0.8 σ) at a resolution of 2.5 Å for the pocket as input (Fig. 2b), the filler ED was generated as shown in Fig. 2c. Interestingly, the generated filler ED replaced the region originally occupied by water molecules and covered the unoccupied cavities (indicated with red arrows in Fig. 2c). Then, the generated filler ED was converted into several reconstructed EDs (Fig. 2d). For each reconstructed ED, a number of map skeletons were generated and then interpreted into fragments. It is not a surprise to observe that the classical hinge binding groups could be well fitted into the reconstructed EDs for a kinase target (Fig. 2e). Then, the fragments were selectively connected to create different molecules (Fig. 2f and 2g).

**Figure 2.**
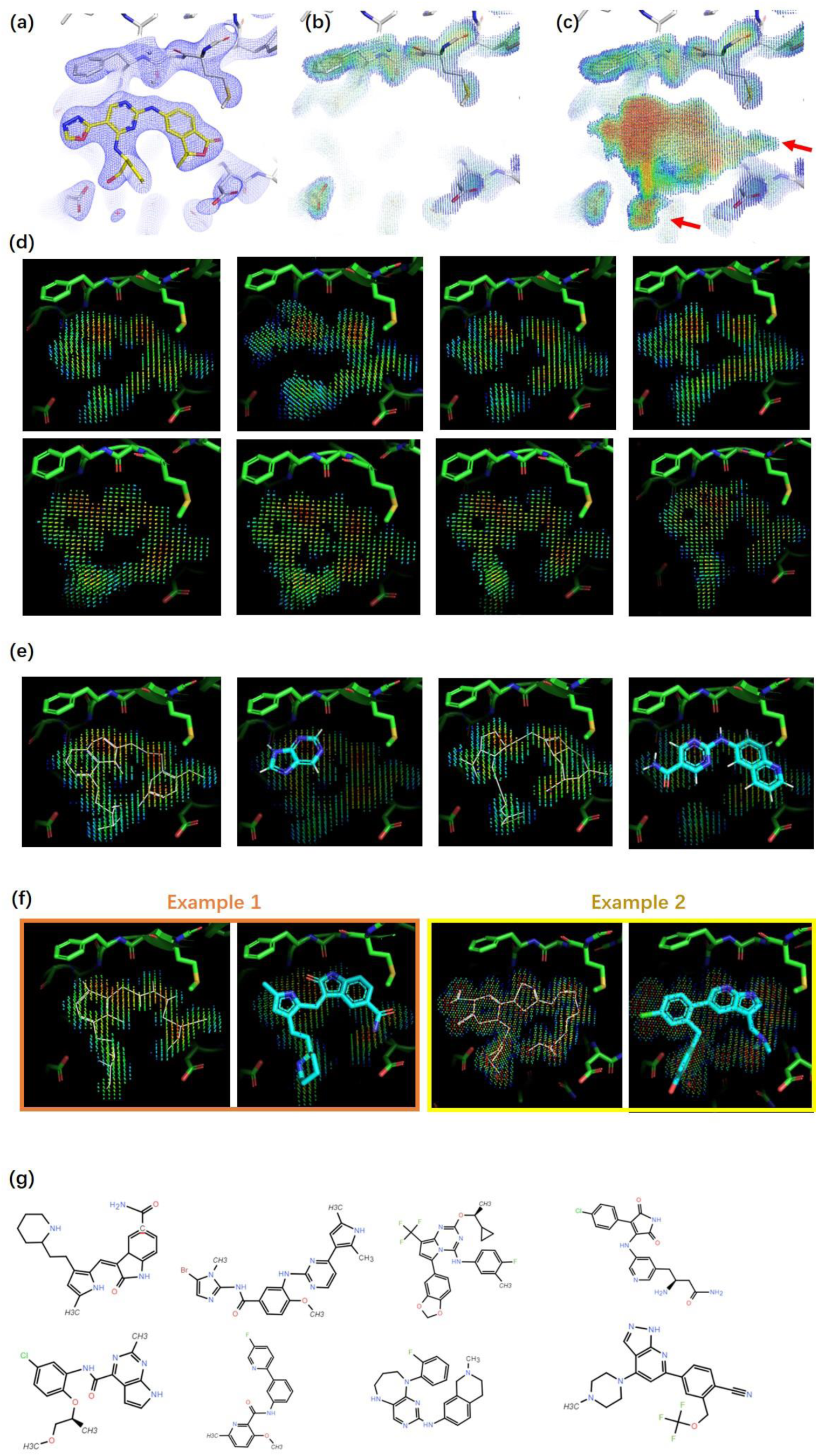
ED based 3D molecule generation for HPK1. a) Binding pocket and reference ligand (PDB code 7KAC). Experimental ED (2Fo-Fc map at 1.2 σ contour level) for the pocket and ligand are shown as a blue mesh. b) Pocket ED (2Fo-Fc map at 0.8 σ contour level) with ligand removed. c) Generated filler ED. For the rainbow color scheme, red indicates a strong ED intensity, and blue indicates a weak ED intensity. The extension of generated ED to the region occupied originally by water molecules and cavities originally unoccupied are indicated by red arrows. d) Reconstructed ED generated based on filler ED. e) Map skeleton and hinge binding fragments interpreted from the map skeleton. Map skeletons are shown as white lines. f) Examples of generated molecules and their map skeleton aligned with reconstructed ED. G) List of examples of generated molecules. ED, electron density.

### Model evaluation

As shown in Fig. 1c, our molecule generative model was evaluated from three perspectives: 1) the ability to generate valid molecules in terms of QED and SAS while maintaining a reasonable diversity; 2) the ability to reproduce classical active compounds, defined as generation of molecules with over 0.5 Tanimoto similarity^23^ with the reference active compounds; 3) the ability to generate novel active compounds, defined as generation of molecules with less than 0.5 Tanimoto similarity with the reference active compounds but possessing a similar binding mode.

We selected three targets including HPK1 (PDB: 7KAC), Covid19-3CL (PDB:7VU6), and VDR (PDB: 1S19), to test the performance of our model. They were selected as representatives of kinase, protease, and nuclear receptors. Regarding references (Table S1), 6334, 1101, and 757 compounds with reported activity were used for HPK1, 3CL, and VDR, respectively (supplementary material 1). In addition, a state-of-art 3D molecule generative model^24^ reported in NeurIPS 2021 was used as a comparison benchmark.

### Validity Test

The most fundamental test for molecules generated by AI models is the validity test evaluating the QED and SAS. We generated 10,000 molecules for each of the three targets using our model and the benchmark model and compared the QED and SAS of the generated molecules. To make the comparison results easy to understand, we also calculated the QED and SAS of the reference compounds and used them as positive controls. As shown in Table 1, although both our model and the benchmark model performed well for QED, our model outperformed the benchmark model on SAS.

**Table 1.** Evaluation of molecular validity.

| Metric for validity test |  | Target |  |  |  |  |  |  |  |  |
| --- | --- | --- | --- | --- | --- | --- | --- | --- | --- | --- |
|  |  | HPK1 |  |  | Covid19-3CL |  |  | VDR |  |  |
|  |  | Ours | BM<sup>a</sup> | Ref.<sup>b</sup> | Ours | BM<sup>a</sup> | Ref.<sup>b</sup> | Ours | BM<sup>a</sup> | Ref.<sup>b</sup> |
| QED | Avg. | 0.54 | 0.55 | 0.47 | 0.40 | 0.62 | 0.32 | 0.55 | 0.48 | 0.49 |
|  | Med. | 0.53 | 0.56 | 0.46 | 0.39 | 0.64 | 0.29 | 0.53 | 0.47 | 0.48 |
| SAS | Avg. | 3.0 | 5.5 | 3.6 | 3.2 | 4.5 | 3.2 | 2.9 | 5.3 | 3.0 |
|  | Med. | 2.9 | 5.5 | 3.6 | 3.1 | 4.5 | 2.8 | 2.9 | 5.3 | 2.8 |

**Notes:** 10,000 molecules are respectively generated by our model and benchmark model for each target; <sup>a</sup>BM refers to a model<sup>24</sup> reported in NeurIPS 2021 and used as benchmark here; <sup>b</sup>Reference compounds (Table S1); About the color scheme: all the values of reference compounds have their cells colored yellow. For generated molecules, if their values are better

### Chemical Space Distribution

To understand the diversity of the molecules generated by our model, we calculated the Tanimoto-similarity-based diversity. As shown in Table S2, the molecules generated by the benchmark model exhibit higher diversity than those generated by our model. However, such general diversity is not suitable for the evaluation of molecules generated with constraints, because well-functioning constraints may reduce the general diversity. To perform a comprehensive evaluation, we referred to the small-molecule-universe representative universal library (SMU-RUL) chemical space^25^, a self-organizing-map (SOM)-supported chemical space used to describe the distribution of molecules with molecular weight less than 500 Da. To simplify the comparison results, samples from PubChem were used to represent molecules lacking target-specific constraints, and active reference compounds are used to represent molecules with tight constraints. As shown in Fig. 3, we observed that our model achieved a better balance between the diversity and constraints than the benchmark model (i.e. the molecule generated by our model is concentrated towards the region where the active compounds are positioned, whereas the molecules of the benchmark model are distracted to some other regions and thus appear more diverse).

**Figure 3.**
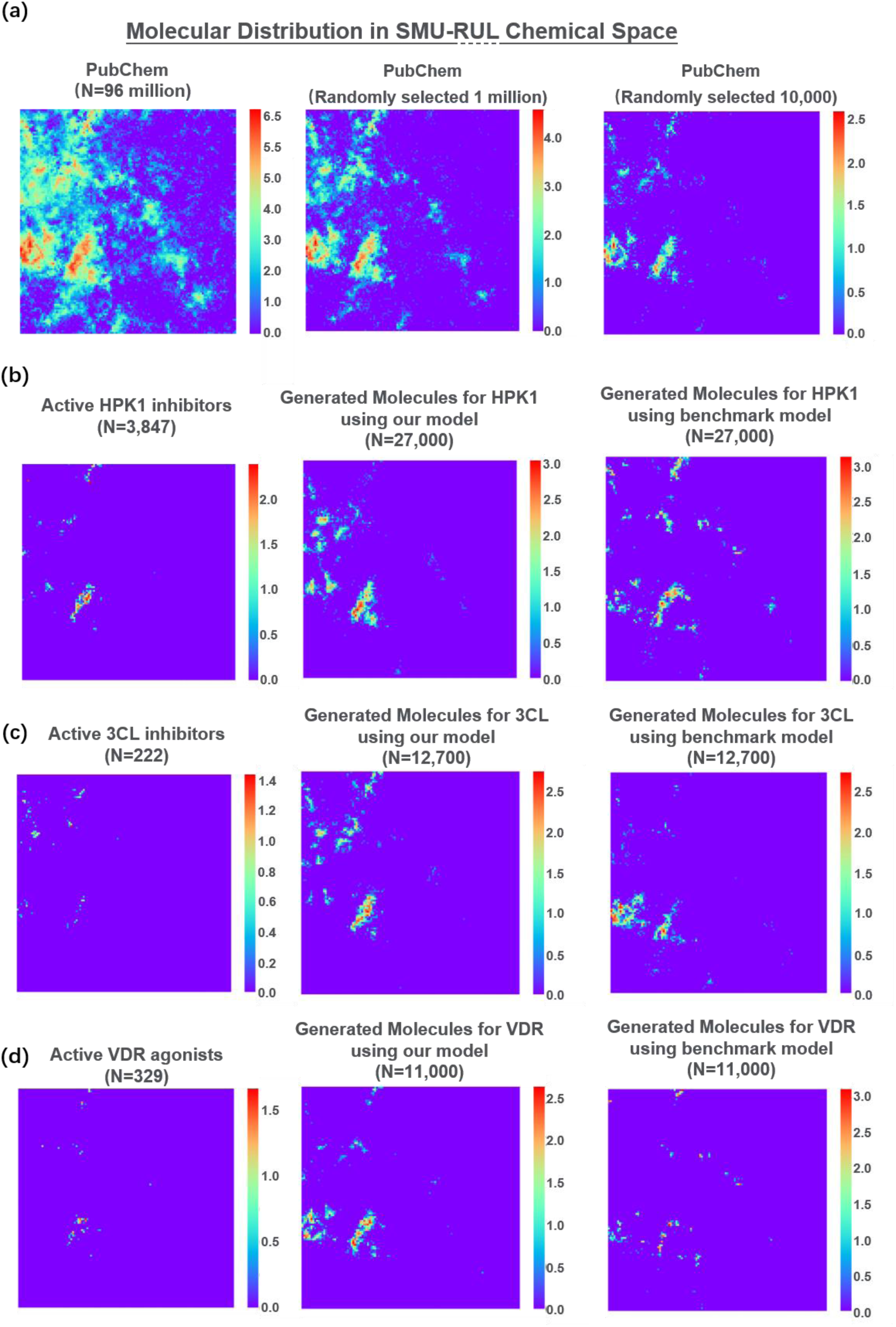
Chemical space distribution of generated molecules and references. A 120 × 120 SOM was created using SMU-RUL^25^ compounds. The color indicates the number of molecules on a logarithmic scale. a) The distribution of PubChem with different numbers of randomly selected samples. b) The distribution of HPK1 reference active compounds and molecules generated using different models. c) The distribution of Covid19-3CL reference active compounds and molecules generated using different models. d) The distribution of VDR reference active compounds and molecules generated using different models. The number of molecules generated by our model is adjusted to match the number that can be generated by the benchmark model within a reasonable time frame. The SMILES of the molecules and their positions in the chemical space are provided in Supplementary material 3. SOM, self-organizing-map; SMU-RUL, small-molecule-universe representative universal library.

### Reproduction of classical active compounds

Besides testing the validity of the generated molecules, we also needed to test whether the molecular generative model can reproduce already known active compounds. To this end, 1 million molecules were generated for each of the three targets and compared to reference compounds by measuring the Tanimoto similarity of the ECFP4 fingerprint. In addition, generating a molecule with over 0.5 Tanimoto similarity against a reference compound was considered a successful reproduction of this reference compound. Our model successfully reproduced reference compounds for all three targets (Supplementary material 2). Taking HPK1 as an example, some reference compounds and their generated counterparts are listed in Fig. 4. Furthermore, when we sorted reference compounds by their activity, we observed that our model tended to generate molecules with higher similarity to active and medium-active compounds, compared with inactive ones, for all three targets (Table 2). Apart from the similarity trend, our model also reproduced more active and medium-active compounds than inactive ones for HPK1 and Covid19-3CL. Regarding the comparison with the benchmark model which failed to generate million-level molecules within a reasonable time frame (Table S3), it would be unfair to use all the 1 million molecules generated by our model. Therefore, our molecules that were subjected to the comparison were randomly selected form the 1 million previously generated to match the capacity of the benchmark model. As shown in Table S4, although neither of the two models reproduced active reference compounds under the condition that only tens of thousands of molecules were generated, our model still provided molecules with better Tanimoto similarity to the references than the benchmark model did.

**Figure 4.**
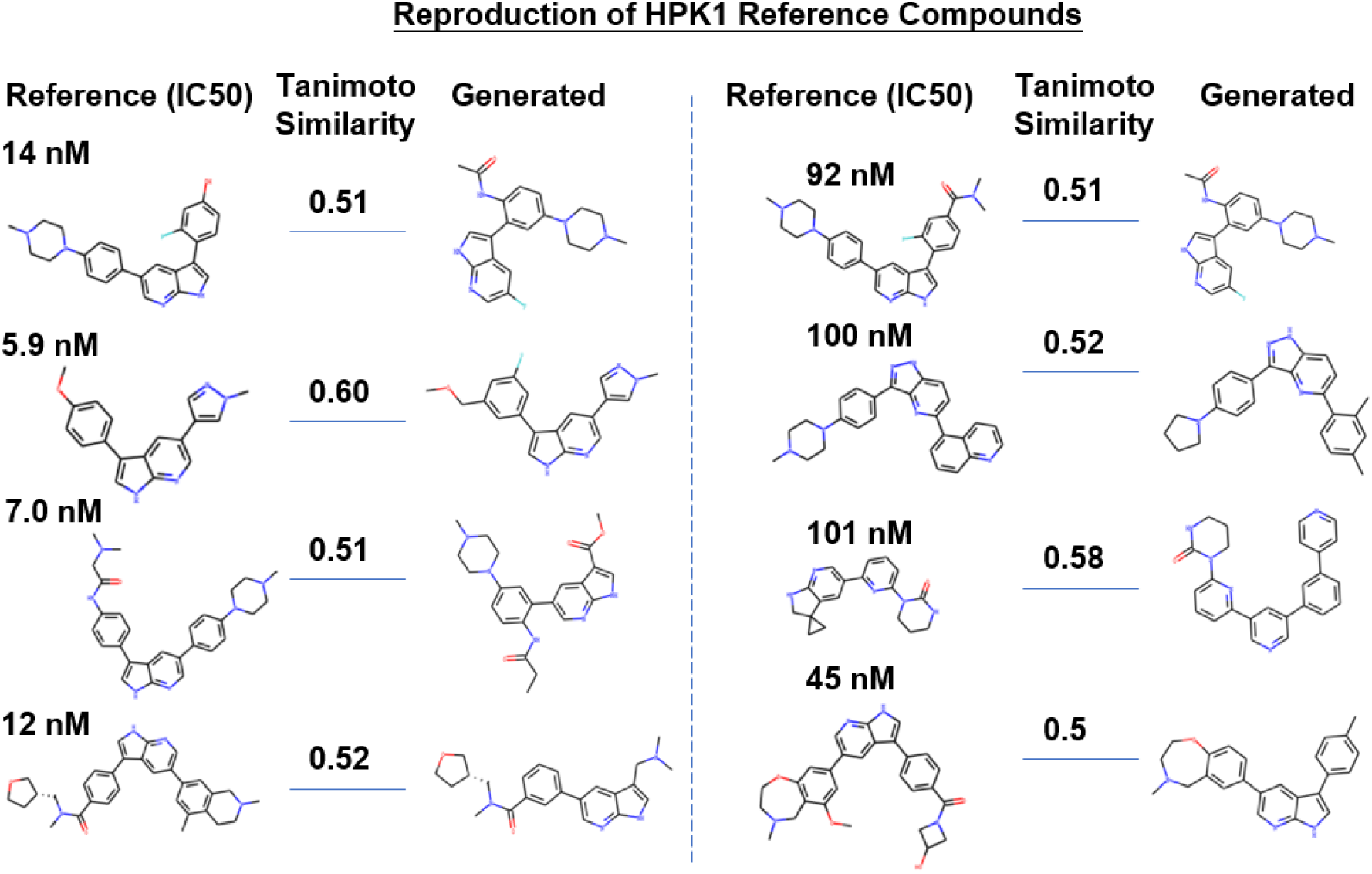
Examples of generated molecules that are similar to reference compounds for HPK1.

**Table 2.** Test of reproducing classical active compounds for HPK1.

| Metric for classical active compounds reproduction test |  | HPK1 |  |  | Covid19-3CL |  |  | VDR |  |  |
| --- | --- | --- | --- | --- | --- | --- | --- | --- | --- | --- |
|  |  | Active | Medium | Not active | Active | Medium | Not active | Active | Medium | Not active |
| # of reference compounds |  | 3847 | 2319 | 168 | 222 | 248 | 631 | 329 | 67 | 361 |
| Our Model | # of molecules generated | 1 million |  |  | 1 million |  |  | 1 million |  |  |
|  | Max. of Tanimoto similarity to ref. cpd. | 0.76 | 0.72 | 0.51 | 0.58 | 0.73 | 0.61 | 0.57 | 0.44 | 0.55 |
|  | # of reference molecule reproduced<sup>a</sup> | 58 | 53 | 1 | 3 | 15 | 11 | 1 | 0 | 7 |
|  | # of generated molecules with over 0.5 Tanimoto similarity to ref. cpd. | 30 | 20 | 2 | 14 | 158 | 42 | 1 | 0 | 10 |

**Note:** <sup>a</sup>reproduced: for a reference compound, if a generated molecule has over 0.5 Tanimoto similarity with it for ECFP4, then this reference compound is considered to have been reproduced. The SMILES of the generated molecules and their similar reference compounds are provided in Supplementary material 2.

### Generation of novel active compounds

If a molecule generative model can only generate classical active compounds, it won’t be attractive to researchers searching for new drugs. To test our model’s ability to generate active compounds with novel structures, we searched the generated library for the molecules exhibiting a <0.5 Tanimoto similarity with reference compounds while sharing a similar binding mode.

Taking HPK1 as an example, 2,769 HPK1 reference compounds with reported Ki or Kd were selected and docked into the 7KAC pocket by using Glide SP^26, 27^ to provide a background for binding mode analysis. This is because the binding mode is more theoretically related to Ki and Kd than to EC50 or IC50. However, as shown in Fig. 5a, the Glide score could not efficiently distinguish the reference compounds with different activities, which implies the need for a more powerful descriptor of binding mode. Thus, we referred to NCI fingerprints depicted using the independent gradient model (IGM) method^28^. The IGM method calculates NCIs by analyzing the topological properties of EDs, and it can provide a full spectrum of NCIs^17^, whereas the traditional rule-based method only provides a short list of classical NCIs. Compound clustering based on the NCI fingerprint was conducted using T-distributed stochastic neighbor embedding (t-SNE)^29^. As shown in Fig. 5b, the clustering results aligned quite well with the activity: compounds sorted into one cluster tended to possess similar activities despite the low discriminative capability for samples with very high activity (separating compounds below 1 nM from those below 10 nM).

**Figure 5.**
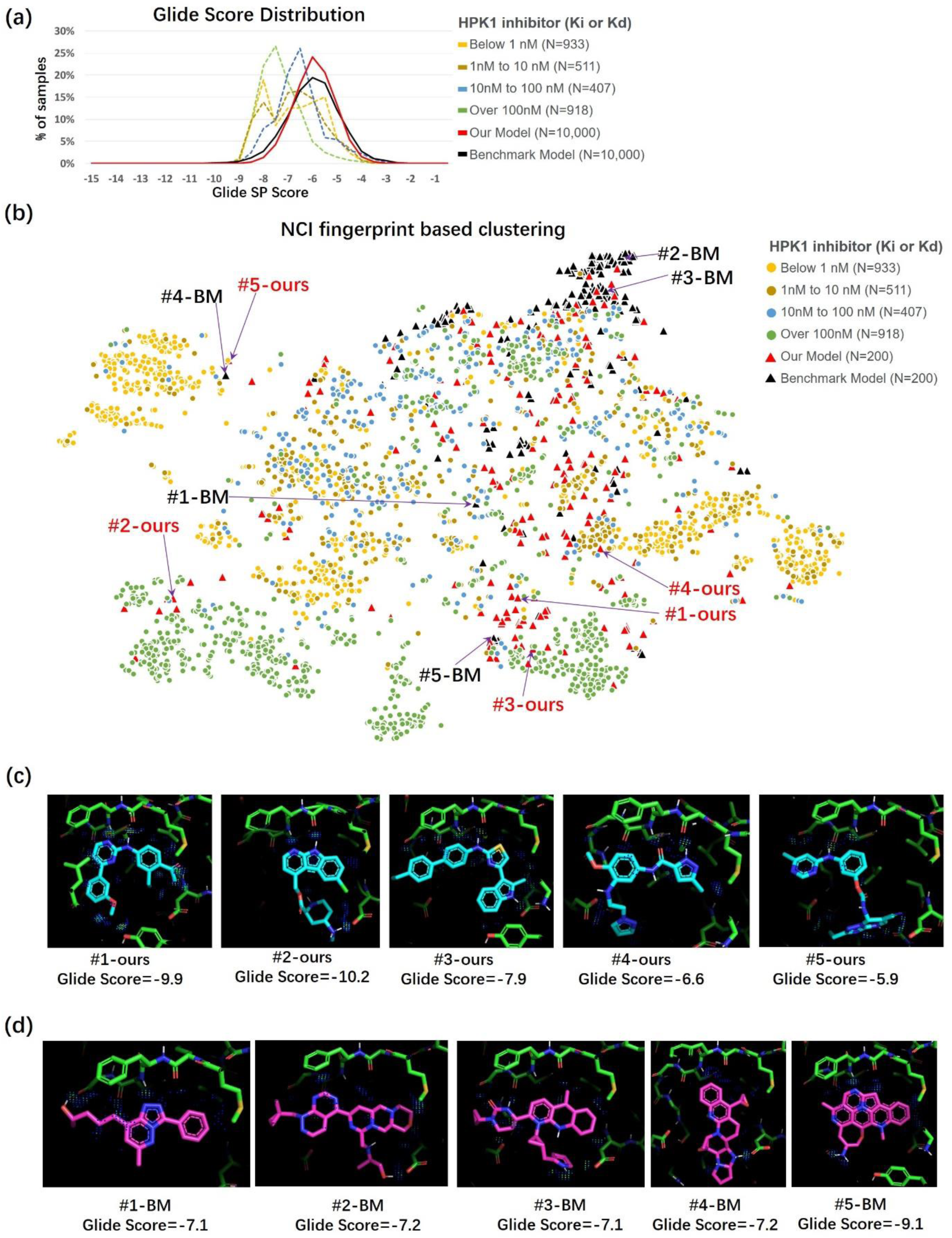
Binding mode analysis of the generated molecules for HPK1. a) Glide Score distribution for active reference compounds and generated molecules. b) Results of t-SNE clustering using IGM calculated NCIs as features. c) Binding mode of selected molecules generated by our model. d) Binding mode of selected molecules generated by benchmark model. For panel c and d, NCI regions are indicated with dots colored using the rainbow scheme, in which blue indicates weak interactions and red indicates strong interactions. Details of docking score, clustering analysis, and NCI fingerprints of selected molecules are provided in supplementary material 4, 5, and 6, respectively. IGM, independent gradient model; NCI, non-covalent interaction; t-SNE, t-distributed stochastic neighbor embedding.

To analyze the binding mode of molecules generated for HPK1, we randomly selected 10,000 generated molecules with <0.5 Tanimoto similarity against the reference compounds and then scored them using Glide by only employing the minimization and scoring functions. This operation was conducted for both our model and the benchmark model. Similarly to the reference compounds, the Glide score was unable to efficiently distinguish between the performances of our model and the benchmark model (Fig. 5a). Therefore, we randomly selected 200 molecules from each of the two models and conducted NCI fingerprint clustering. As shown in Fig. 5b, the molecules generated by the benchmark model tended to be more concentrated and farther from the cluster of active compounds than those generated by our model. Molecules at different positions on the NCI fingerprint map are listed in Fig. 5c and 5d, and their NCI fingerprints are provided in Fig. S1. The hinge region-related NCIs, which are considered crucial for active compounds, were exhibited for molecules generated close to the cluster of active compounds. These NCIs were weak for molecules far from the clusters, such as #2-BM and #3-BM. Although there were cases (#4-BM and #5-ours) where the molecule from our model was close to the benchmark model molecule in the NCI fingerprint map, and both of them were close to the cluster of active compounds, our molecules are still easily distinguished for their superior synthetic accessibility.

## Discussion

To solve the fundamental problem of target compound identification for structure-based drug design, we used experimental ED to train an AI 3D molecule-generative model for the first time. The core of our approach can be understood from three perspectives: data, molecule representation, and model architecture. Since experimental EDs contain experimentally observed time-averaged conformational change and solvent information that are critical for ligand binding, the introduction of these ED data to AI molecular generation provides a promising solution to the challenge of insufficient data. In addition, we highlighted the value of representing molecules as ED; such representations are not only compatible with the usage of experimental ED data during model training but also avoid the data sparsity problem caused by using multiple channels. Furthermore, we proposed a procedure allowing for step-by-step constraint learning, just as structural biologists build small molecules in the pocket. In summary, we presented a novel solution for AI-based 3D molecule generation.

One possible limitation to our approach is associated with a scenario in which the model is used without experimental ED data as input. Such a scenario could occur when the generative model is used for a pocket provided by a molecular dynamic approach. For our model, although it is trained using experimental ED data as input, it also works when using a calculated Fourier-synthesis based ED map as input. Specifically, one just needs to add reasonable B factors (e.g., 20 with a Gaussian perturbation) for each atom and then calculate the ED map using the Fourier-synthesis.

To further improve the performance of our model, we are considering the use of multi resolution ED. For the training of our current model, all pockets were represented as ED at a resolution of 2.5 Å, and therefore, some of the PDB entries with a resolution higher than 2.5 Å had their data at high resolution shells unused. One possible method for utilizing these valuable data is to create multiple channels for different resolutions. However, one must strike a balance between creating multiple channels and avoiding the data sparsity problem for high resolution channels receiving PDB entries with low resolution data.

### Methods

### 1. Molecule Training

We designed a two-step generation scheme using ED to represent molecules. For the first step, we used 27,006 complex data to focus on the learning of the complementarity between protein pockets and ligand molecules. For the second step, we trained the model to generate valid molecules under constraints contained in the generated ED in the first step. Decoupling the two steps makes it possible to use millions of QM and force-field-generated computational data for model training in the second step. ED was featured using 3D CNN, and data augmentation was used to achieve rotational invariance.

#### 1.1 Ligand ED Generative Model

Similar to the image generative model (such as pix2pix^30^) used for image inpainting, our model was designed to generate ligand ED (also referred as filler ED) from pocket ED while maintaining complementarity in terms of shape and NCI. The model was built on the GAN framework with the architecture shown in Fig. 1a.

To prepare data for model training, the experimental EDs for pockets were generated using Phenix^31^ and experimental ED coefficients downloaded from PDB. These pocket EDs were used as input features for the AI model. The computational ED for ligands were prepared as labels. The software xtb^32^ were used to calculate computational ED for ligands at GNF2-xTB level using ligand coordinates as input. Since the ED intensity within the nuclear region is much stronger than that within the region where the NCIs take place and where the shape of the molecule is defined (i.e., the isosurface at 0.03 e/Å^3^), the range of values were compressed using logarithms and then used as labels for the AI model.

The pocket EDs were input into a V-Net 3D image generation network, which acts as a generator. In addition to the generator, two discriminators, called complex and ligand discriminators, were designed. The complex discriminator was responsible for examining the level of complementarity between the ligand and the pocket, and the ligand discriminator was responsible for verifying ED validity from the perspectives of gradient and connectivity.

The protein-ligand pairs used to train the filler ED generative model were from the PDBbind database^33^. In total, 27,006 ligand-protein pairs were extracted from 12,905 complexes. The number of the ligand-protein pairs was larger than that of the complexes because we included drug-like molecules as well as other binders with molecular weight less than 600, such as ATP, ADP, and sugars. To avoid the appearance of similar pockets in both the training and the testing sets, training-testing split was done by referring to the pocket classification from previous studies focusing on the 1D and 3D pocket similarity^11, 34^. The key NCIs identified using a previously reported method^17^ were also included in the model during training with the purpose of making the network more attentive on the NCI related regions. Smooth L1 loss was performed separately to strengthen the loss of the NCI related regions.

The GAN and its loss were expressed using the following equations:

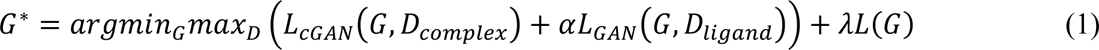

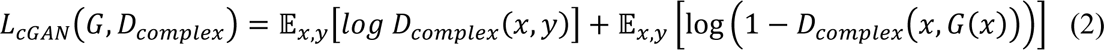

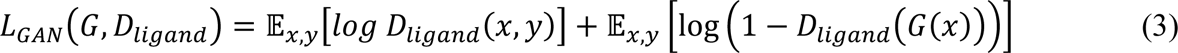

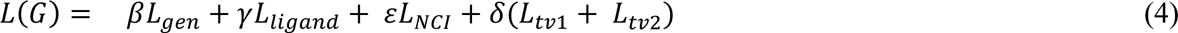

where *D_complex_* represents the complex discriminator, and *D_ligand_* represents the ligand discriminator. *L(G)* indicates the regression loss of GAN, *x* represents the input pocket*, y* represents the ground truth ligand, and *α, λ, β, γ, ε,* 𝛿 are parameters to be learned. *L_tv1_* and *L_tv2_* are used to measure the similarity of two EDs from the perspectives of the first and second orders derivative of intensity, respectively.

The components of *L(G)* are defined as follows:

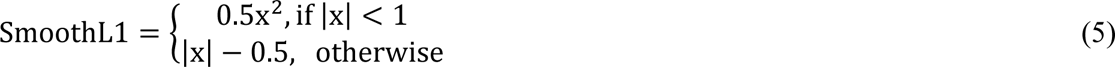

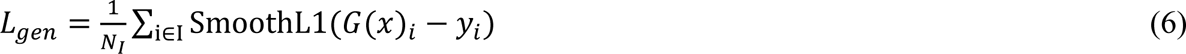

where *I* indicates the entire region, and *N_I_* indicates the number of elements in 𝐼.

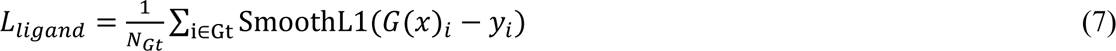

where *Gt* indicates the region covered by the ground truth, and *N_Gt_* indicates the number of elements in *Gt*

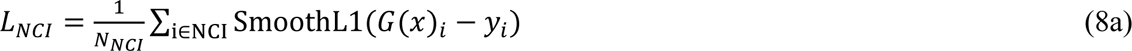

where *NCI* indicates the region covered by *NCI*, and *N_NCI_* indicates number of elements in *NCI*

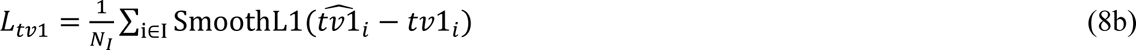

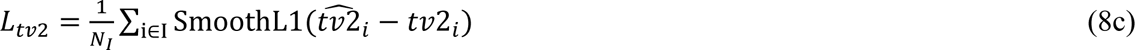

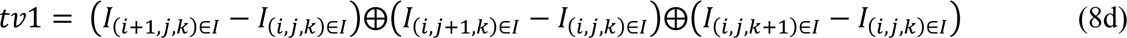

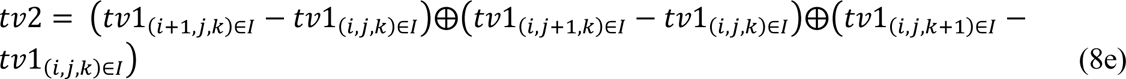

where ⊕ indicates the operator of concatenate.

#### 1.2 Ligand Structure Generative Model

The model was designed to infer the atom type and coordinates of small molecules using generated filler ED as input. A PixelCNN was trained to do the sampling with the generated filler ED as a condition in the latent space constructed using a VQ-VAE2 framework. A group of reconstructed EDs that are more similar to a molecule were generated in the sampling process. Next, the reconstructed ED was submitted to a V-Net framework for the detection of key atoms. This step is similar to that of key point detection in human skeletons in the field of image processing but differs in that it faces severe complications associated with the flexible number of points, diverse types of points, and complicated topology.

VQ-VAE2 was used to compress the ED into a discrete latent space and learn a codebook in which EDs are represented by a series of embeddings. Each pixel in the latent space can be described by an integer in the codebook and the integer can be logically considered as a 3D molecular fragment. To retain the constraints in the input ED while ensuring that the output ED can match a valid molecule, a separate autoregressive prior (PixelCNN) was taught to sample the latent space with input ED as a given condition. Two codebooks were learned during training: the top-level codebook focusing on the extraction of general profile information (such as shape) and the bottom-level codebook focusing on the detailed information (such as local conformations). Regarding sampling with the two codebooks, the input ED was used as a condition to sample the top-level codebook, and the top-level encoding was used as the condition to sample the bottom-level codebook. Thus, the relationship between the conditional ED (i.e., input ED) and the generated ED was decoupled.

The input ED value at each pixel was normalized to a range between -1 and 1 using tanh plus batch normalization in the encoder before it proceeded to the codebook for sampling. This improves the codebook encoding utilization, and thereby increases the amount of information represented through encoding.

The data used to train the VQVAE2-PixelCNN-VNet model for inferring atoms from generated ligand ED were obtained from PubChem. Two million molecules with a QED exceeding 0.3 were extracted. For each molecule, 20 conformers were generated using ConfGen^35^. EDs of the 40 million conformers were generated using xtb^32^ and then used as training data for the VQ-VAE2. The V-Net training, which was used to infer the atom location, was trained using the above 40 million EDs as features and the conformers used to generate these EDs as labels. The V-Net output was comprised of connected atoms and called a map skeleton.

EDs were voxelized into a discrete 0.5Å cubic grid with a side size of 24Å. There were represented as a tensor p ∈ ℛ^1×48×48×48^, where channel size was 1, expressing the intensity of ED.

The VQ-VAE2 loss is shown in Eq (9),where *x* is the training instance, *D* is the decoder of the VQ-VAE2 and *e* is the encoder. The reconstruction loss is shown in Eq (10). It combines the L_SoftDiceLoss_ as well as L_SmoothL1_ . To generate better ED reconstructed shapes, we adopted L_SoftDiceLoss_ as shown in Eq (11) for segmenting ED and the background where *y* is the target and ŷ is the prediction. The target label of the pixel with the ED value was set to 1, and the rest of the background part was set to 0. Further, L_SmoothL1_ was obtained using Eq (12); similarly, y is the target and ŷ is the prediction.

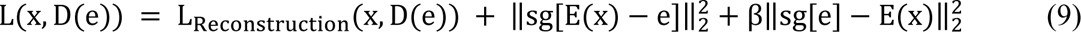

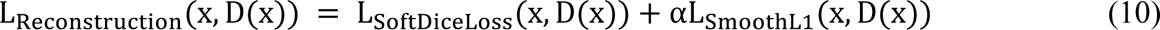

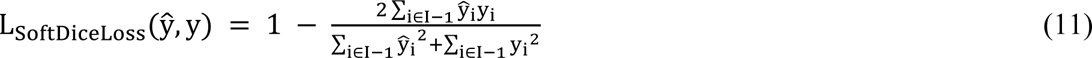

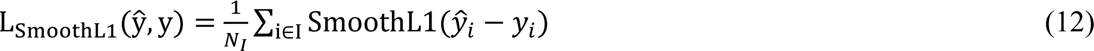

*PixelCNN* optimized the negative log-likelihood of the training data *to maximize the probability* 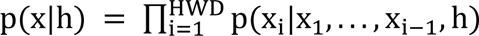, where H represents height, W represents width, and D represents depth of the 3D cube where h is the condition. Top-level and bottom-level prior networks were modeled with 6×6×6 and 12×12×12 latent variables, respectively. Additionally, the condition of the top-level PixelCNN was the previously generated filler ED. The bottom-level network was conditioned on the top-level prior.

Respecting the V-Net output, atoms from molecules were characterized as one-hot representations in each cell of the 3D grid. The ground truth of the atom types was represented as a tensor, where 10 channels were denoted as the PAD (no atoms), C, N, O, S, P, F, Cl, Br, and I, respectively, in this work. Then, V-Net utilized the cross-entropy loss to classify atom types in each voxel x ∈ ℛ^3^.

#### 1.3 Map skeleton substitution and fragment assembling

The V-Net generated map skeletons that were already positioned in the pocket. A map skeleton was then divided into two fragments, and each fragment was replaced by molecular fragments with similar shapes from the conformation library. Briefly, the library entries with a root mean square distance less than 1.5 Å from the fragment of the map skeleton were used to substitute that fragment.

The molecules from PubChem were matched with 35 reaction templates^36^ according to the substructure superposition supported by RDKit^37^, and then cut into two fragments at the matching site. If a molecule matched multiple reaction templates, it was still cut into two fragments each time, but the cutting site was different. In this way, over 270,000 fragments with labeled cutting sites were obtained. Then OpenBabel^38^ was used to generate a maximum of 20 conformations for each fragment, and it finally produced a library with 3.5 million 3D conformations.

The map skeleton was cut into several fragment pairs in the following manner: each time the whole molecule was cleaved at one acyclic single bond, two fragments were generated; next time, the whole molecule was cleaved at another acyclic single bond; the process was repeated until all acyclic single bonds had been cut. For every fragment pair, the above-established 3.5 million 3D conformation library was searched for entries that were similar to the fragments. Next, the two groups of selected entries were assembled together in an enumerative way to make a list of target molecules. Subsequently, several filters were applied to remove the unqualified molecules. These filters include the following:

1. Collides with pocket: the distance from the pocket heavy atom is less than 2.5 Å.
2. Collides between the fragments: the distance from the non-bonded heavy atom is less than 2.0 Å.
3. Stability and synthetic accessibility filters and drug-like filter reported by Virshup et al.^25^

### 2. Model Evaluation

The data we used for chemical space construction consisted of 8.8 million molecules from SMU-RUL^25^.

The evaluation data were collected from 144 publications including research papers and patents.

### 3. Chemical space construction

Chemical space was constructed mainly based on the method described in a previous study^25^. Specifically, Moreau−Broto autocorrelation descriptors were used for molecule description. These descriptors encode topological correlations between the atomic properties of a molecule into a fixed-length vector.

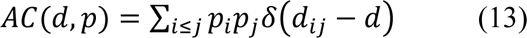

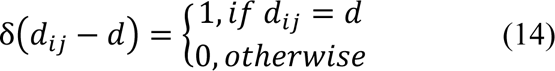

where d*_ij_* is the number of bonds separating atoms *i* and *j*, while p*_i_* is the value of atomic property *p* on atom *i*. In our work, *p* includes four properties: carbon-scaled atomic mass, carbon-scaled atomic van der Waals volume, carbon-scaled atomic Sanderson electronegativity, and carbon-scaled atomic polarizability. The values of *d* ranged from 0 to 7. As a result, a molecule was finally described as a vector with 32 dimensions. The autocorrelation descriptors were calculated using PyBioMed^39^.

Up to 8.8 million molecules from SMU-RUL^25^ were converted into 32D vectors and used for the construction of the chemical space by using the SOM method^40^. The SOM size was 120 ×120. The SOM was implemented using MiniSom library^41^.

### 4. NCI fingerprint analysis

Docking was implemented by using Glide^26, 27^. IGM-based NCI analysis was implemented using Multiwfn^42^. The atom-pair-based NCI list output by Multwfn was further annotated with the Mol2/Sybyl atom types for both of the atoms using OpenBabel^43^ and PyBel^38^ packages, and then submitted to Scikit-learn^44^ for t-SNE^29^ clustering analysis.

### 5. Benchmark Model

The source code of benchmark model is downloaded from its official website: https://github.com/luost26/3D-Generative-SBDD

### 6. Software for Figures and Tables

The structure and ED figures were made using Pymol.

## Supporting Information.

SI_1_Reference_Compounds.csv

SI_2_Table 2 and Figure 4 Data Generated_Molecules_Similar_to_Ref.csv

SI_3_Figure 3 Data Chemical_Space_Projection.csv

SI_4_Figure5a_Glide Score.csv

SI_5_Figure5b_NCI FP Clustering.csv

SI_6_Figure5c NCI_FP.csv

## AUTHOR INFORMATION

Author Contributions

Bo Huang and Wenbiao Zhou conceived the idea. Bo Huang provided instructions for experimental ED processing, molecular representations and model evaluation. Wenbiao Zhou provided instructions on AI models. Lvwei Wang developed the GAN module. Rong Bai developed the VQVAE2-PixelCNN-VNet module. Xiaoxuan Shi developed the fragment assembling code and conducted the evaluation. Wei Zhang and Yinuo Cui prepared evaluation data. Cheng Wang supported chemical space distribution analysis. Haoyu Chang supported GAN training. Xiaoman Wang supported website development. Yingsheng Zhang and Jielong Zhou provided instructions on general model architecture. Wei Peng provided instructions on the selection of validation targets.

## DATA AVAILABILITY

Reference compounds extracted from literatures, data used for chemical space distribution analysis and NCI fingerprint analysis, and results of reference compound reproduction are available as supplementary materials.

## CODE AVAILABILITY

Partial codes are available upon request.

Our model provides service to academic users via https://edmg.stonewise.cn/#/create

## COMPETING INTERESTS

The authors declare no competing interests.

## ACKNOWLEDGMENT

All the authors received funding from StoneWise Ltd.; B.H. and W.Z. also received funding for cloud computing from Beijing Municipal Science & Technology Commission project (No. Z211100003521001).

---

**Figure S1.**
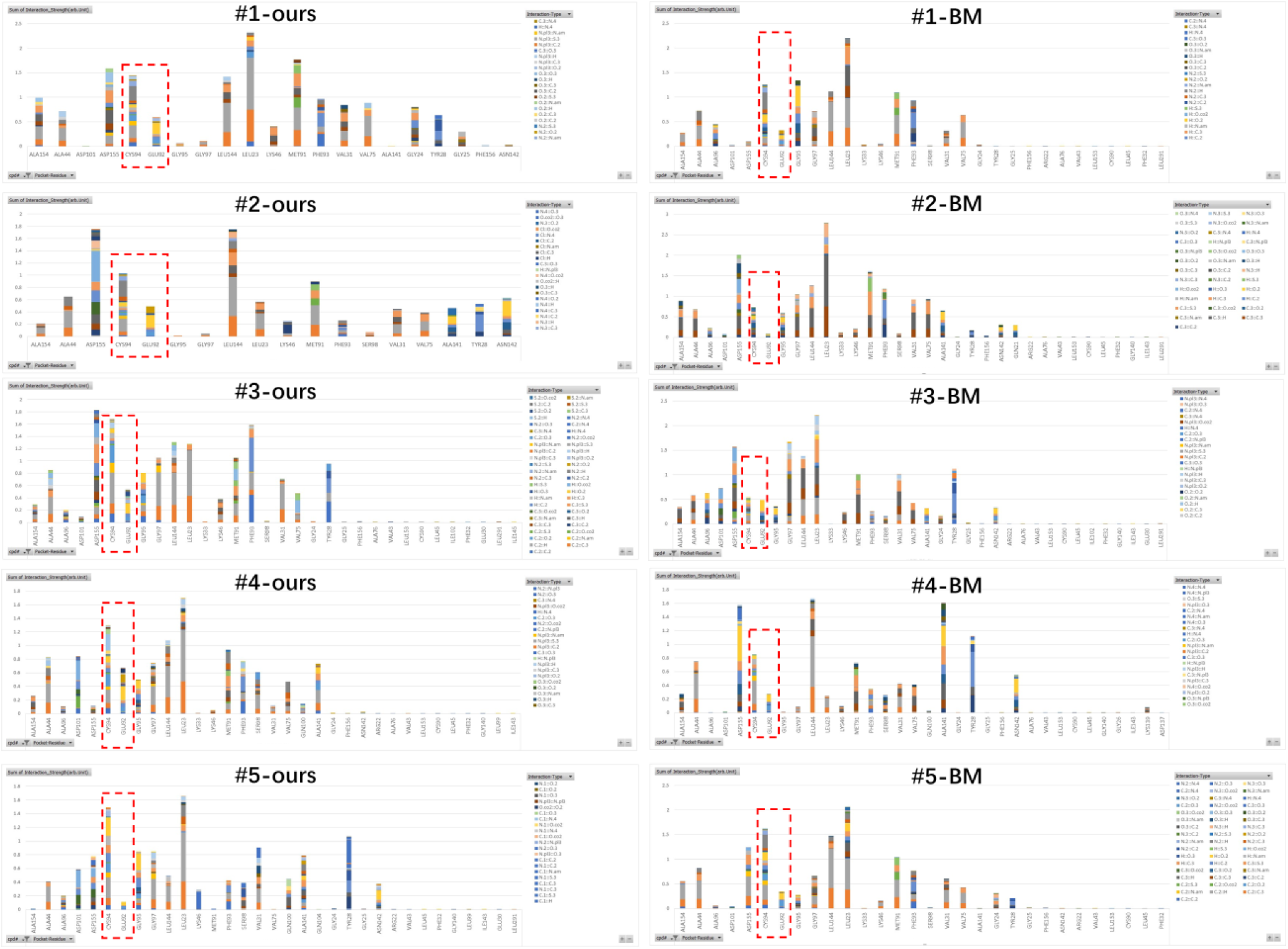
NCI fingerprint of selected molecules. Hinge region related NCIs (i.e. E92 or C94 involved) are indicated with red box. More details are provided in supplementary material 6.

**Table S1.**
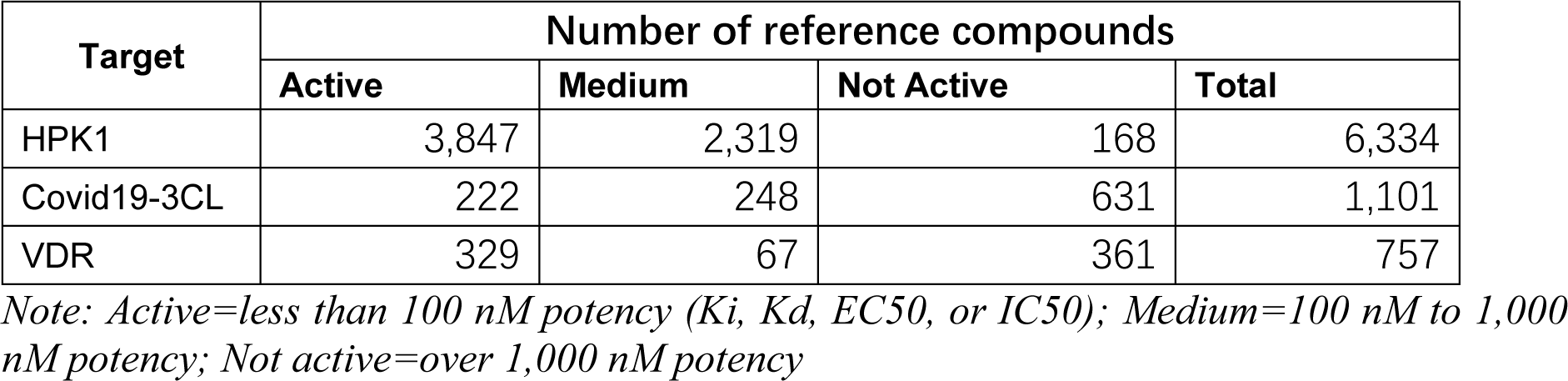
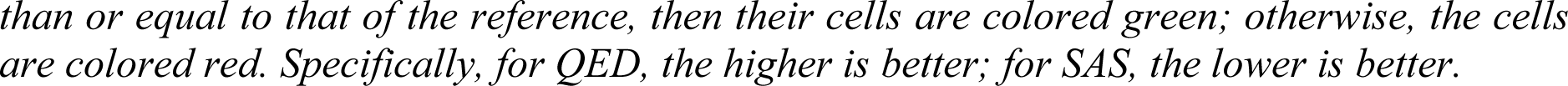
Reference Compounds.

**Table S2.** Diversity of molecules generated by our model.

| Diversity |  | Target |  |  |  |  |  |  |  |  |
| --- | --- | --- | --- | --- | --- | --- | --- | --- | --- | --- |
|  |  | HPK1 |  |  | 3CL |  |  | VDR |  |  |
|  |  | Ours | BM<sup>a</sup> | Ref.<sup>b</sup> | Ours | BM<sup>a</sup> | Ref.<sup>b</sup> | Ours | BM<sup>a</sup> | Ref.<sup>b</sup> |
| Diversity<sup>a</sup> | Avg. | 0.86 | 0.89 | 0.82 | 0.86 | 0.90 | 0.68 | 0.86 | 0.86 | 0.64 |
|  | Med. | 0.87 | 0.90 | 0.85 | 0.86 | 0.91 | 0.84 | 0.87 | 0.86 | 0.54 |

**Note:** <sup>a</sup>Diversity=1- (the average of pairwise Tanimoto similarities over ECFP4 fingerprints among the molecules). The number of reference molecules subjected to the analysis is the same as that of Table 1. For each target, 10,000 molecules are respectively generated and subjected to the analysis.

**Table S3.** Speed Comparison.

|  | Number of Unique Molecules Generated in 12 hours* |  |  |
| --- | --- | --- | --- |
|  | HPK1 | Covid19-3CL | VDR |
| Our Model | 1,142,718 | 265,152 | 560,328 |
| Benchmark Model | 2,395 | 4,170 | 1,965 |

Note: \*With one NVIDIA® GeForce® RTX 2080 Ti

**Table S4.** Comparison with Benchmark Model in Reproducing Reference Compounds.

| Metric for classical active compounds reproduction test |  | HPK1 |  |  | Covid19-3CL |  |  | VDR |  |  |
| --- | --- | --- | --- | --- | --- | --- | --- | --- | --- | --- |
|  |  | Active | Medium | Not active | Active | Medium | Not active | Active | Medium | Not active |
| # of reference compounds |  | 3847 | 2319 | 168 | 222 | 248 | 631 | 329 | 67 | 361 |
| Our Model | # of molecules generated | 27,000 |  |  | 12,700 |  |  | 11,000 |  |  |
|  | Max. of Tanimoto similarity to ref. cpd. | 0.49 | 0.47 | 0.44 | 0.38 | 0.50 | 0.43 | 0.42 | 0.42 | 0.50 |
| Benchmark Model | # of molecules generated | 27,000 |  |  | 12,700 |  |  | 11,000 |  |  |
|  | Max. of Tanimoto similarity to ref. cpd. | 0.36 | 0.37 | 0.35 | 0.34 | 0.47 | 0.52 | 0.31 | 0.33 | 0.4 |

## Notes

https://edmg.stonewise.cn/#/create

### Competing Interest Statement

The authors have declared no competing interest.

### Summary of Updates

Table S4 has been added.

